# The language network does not support music perception across a range of music experience

**DOI:** 10.64898/2026.09.03.749155

**Authors:** Halie A. Olson, Xuanyi Jessica Chen, Dana Boebinger, Hilary Richardson, Evelina Fedorenko

## Abstract

Music and language share many similarities, yet music does not elicit a response in the language brain areas in adults with typical amounts of music experience, and patients with aphasia show preserved music ability. However, this dissociation might not generalize across development or expertise: it may not yet be present in children, who have limited experience with language and music, or it may be diminished in musicians with extensive music training. To evaluate these possibilities, we here used a precision fMRI approach to examine the responses in the language areas of children (aged 4-16 years, n=80) and adult musicians (n=10) to music. We found that the results obtained previously for non-musician adults replicate in two new control samples and, critically, generalize to both of the newly tested populations: the language areas are already selective for language relative to music in young children, showing no age-related change from early to late childhood, and remain selective despite extensive music training in highly experienced musicians. Thus, it appears that the amount of music exposure and practice have little effect on how the language brain areas respond to music.

**SIGNIFICANCE STATEMENT:** For centuries, philosophers, linguists, psychologists, and neuroscientists have noted similarities between language and music, leading to the hypothesis that these two domains may share processing mechanisms in the human brain. This hypothesis has not found empirical support: in adults with typical amounts of musical experience, language and music processing appear to draw on distinct systems. Here we asked whether the amount of experience with music affects this dissociation. We tested both children, who have less music and language experience than a typical adult, and highly trained musicians with extensive music experience, and found that the language system does not respond to music in either of these populations. These findings suggest that selectivity for language is not affected by music experience.

## INTRODUCTION

Language and music are generative combinatorial systems that rely on hierarchical structures (Fay, 1971; Roads & Wieneke, 1979; Sundberg & Lindblom, 1976; Swain, 1995) and are canonically auditory. Linguistic ability and sensitivity to music structure emerge early in human development (Kuhl, 2000; Trehub, 2003; Winkler et al., 2009; Zentner & Eerola, 2010) and appear to be ubiquitous across human societies (Mehr et al., 2019). These and other similarities led to the hypothesis that language and music draw on overlapping brain mechanisms (e.g., (Koelsch & Friederici, 2003; Patel, 2003, 2008; Tillmann, 2012)). Indeed, some behavioral (e.g., (Fedorenko et al., 2009; Slevc et al., 2009)) and neuroimaging (e.g., (Levitin & Menon, 2003; Maess et al., 2001; Musso et al., 2015; Sammler et al., 2009)) studies have reported evidence for overlap. However, dissociations in patients with brain damage challenge this idea. Some patients with severe aphasia retain music abilities (Basso & Capitani, 1985; Chen et al., 2023; Luria et al., 1965; Slevc et al., 2016), and some patients show music deficits alongside intact language (Peretz et al., 1994, 1997; Piccirilli et al., 2000). Furthermore, fMRI studies using within-individual comparisons and ruling out overlap due to shared attentional resources have found no response to music in the language brain regions (Chen et al., 2023; Deen et al., 2015; Fedorenko et al., 2011; Rogalsky et al., 2011; Sueoka et al., 2024). Indeed, the language areas, which support word retrieval and combinatorial processing in comprehension and production exhibit strong selectivity for language relative not only to music, but many other non-linguistic domains (for review, see (Fedorenko et al., 2024)). In contrast, brain areas that respond to music – located in bilateral temporal cortex, outside primary auditory areas (Binder et al., 2000; Fedorenko et al., 2012; S. Norman-Haignere et al., 2015; Patterson et al., 2002) – are selective for music relative to other types of sounds, including speech (S. Norman-Haignere et al., 2015; S. V. Norman-Haignere et al., 2022).

One limitation is that most past studies focused on adults with average amounts of music experience. Might the neural infrastructure for language and music differ earlier in life, when individuals have less experience with both domains, or in individuals with extensive music training? In **development**, human infants are typically exposed to speech and music in utero, which the auditory system can detect by the third trimester (Birnholz & Benacerraf, 1983). Early-emerging auditory mechanisms may initially process both kinds of inputs and segregate later in life, with experience. Furthermore, the domain-general ability to learn regularities in auditory inputs (Saffran, 2003; Saffran et al., 1996) may support both language and music development (Daikoku, 2018; Mandikal Vasuki et al., 2016; Saffran & Kirkham, 2018), which could rely on overlapping brain areas in childhood. Finally, both speech and music facilitate bonding with caregivers (Savage et al., 2021), which may also lead to neural overlap. These mechanisms would predict higher responses to music in language brain regions in children.

What about **extensive music experience**? The effects of expertise on brain organization have been investigated across domains, from spatial memory in taxi drivers (Maguire et al., 2000), to olfaction in sommeliers (Banks et al., 2016; Filiz et al., 2022), to language in simultaneous interpreters (Hervais-Adelman & Babcock, 2020; Rinne et al., 2000) and polyglots (Jouravlev et al., 2021; Malik-Moraleda et al., 2024). In some cases, expansion of the implicated brain areas has been reported (Banks et al., 2016; Filiz et al., 2022; Maguire et al., 2000), but in other cases, less spatially extensive responses have been observed (Jouravlev et al., 2021; Malik-Moraleda et al., 2024). For musicians, Boebinger et al. (2021) did not find differences in the size and selectivity of music-responsive brain areas relative to non-musicians. However, they did not specifically probe responses to music in the language areas, which might prove more sensitive for detecting differences outside the music areas.

To evaluate the role of music experience on how the language network responds to music, we conducted analyses on two existing datasets: one included children aged 4-16 years, along with adult controls, and the other included adult musicians and non-musician controls. To foreshadow our key findings: the language areas show selectivity for language, with no engagement during music perception, in both children and musicians, thus generalizing earlier results from adults with typical amounts of music exposure across the spectrum of music experience.

## MATERIALS & METHODS

### Participants, design, and procedures

#### Study 1 (Children)

##### Participants

80 children (ages 4.56-16.24 years; mean(SD)= 8.83(2.24) years; 29 female; 65 right-handed/8 left-handed/1 ambidextrous/6 *NA*) and 24 adults (ages 18-59 years; 13 female; 24 right-handed) were included in Study 1. A subset of these participants (n=32) were scanned while wearing a light-exclusion blindfold for the purposes of a different study (Bedny et al., 2015). All participants were native English speakers and had at least 2 usable runs of the fMRI task (with less than 40% of the total volumes excluded as outliers) to be included in these analyses. (Several publications have used different subsets of these data to address distinct research questions (Bedny et al., 2015; Gweon et al., 2012; Ozernov-Palchik et al., 2026; Richardson et al., 2020, 2023).)

##### fMRI task

The fMRI task used a long-event-related design with three conditions critical for the present study: language, foreign speech, and music. The language blocks consisted of 20-second engaging stories recorded by female native English speakers. The stories varied in content and were subdivided into three conditions: mental (stories about characters’ mental states), social (stories about characters’ appearances and relationships), and physical (stories about physical objects and events); for the purposes of the present study, we collapsed across these sub-conditions. The foreign speech blocks consisted of 20-second stories recorded by female native speakers of languages unfamiliar to the participants (Hebrew, Korean, and Russian). The music blocks consisted of 20-second instrumental music clips (piano, violin, guitar, and saxophone). Across conditions, each story/music clip was followed by an auditorily presented question (“Does this come next?”; 1.5 s in duration), and then by another short (3 s) probe clip. The probe clip was followed by a 6.5 s pause during which participants were instructed to respond by pushing one of two buttons (“Yes” or “No”) on an in-scanner button box. Correct responses were followed by an encouraging phrase (“Way to go!”), and incorrect responses by “Let’s try another!”. For the language stimuli, half of the stories were followed by the correct ending (and thus required the “Yes” response); incorrect endings were drawn randomly from all other English story conditions. For the foreign speech stimuli, half of the stories were followed by a probe clip in the same language (and thus required the “Yes” response), and the other half by a probe clip in another foreign language. For the music stimuli, half of the music clips were followed by a probe clip with the same instrument (“Yes” response) and the other half by a probe clip with a different instrument. Participants completed up to four runs, each consisting of 2 experimental events per condition (a total of 6 events for the language condition, per run) and three rest periods 12 s each), for a total run duration of 6.6 min. A rest period occurred at the start of each run, at the halfway point of each run (after five experimental events), and at the end of each run. Condition order was palindromic and varied across runs. Task materials are available on OSF (https://osf.io/cbw6f/). During the fMRI scanning session, child participants were monitored by an experimenter in the control room and a second experimenter who stood next to the scanner bore. For young children, the second experimenter monitored motion and would place a hand on the child’s leg if they were moving noticeably as a reminder to stay still. Participants wore MR-compatible earphones in the scanner and were provided with an MR-compatible button box to respond to task prompts.

#### Study 2 (Musicians)

##### Participants

10 musicians (mean(SD) = 23.5(3.3) years of age; 8 female), all of whom began musical training before 7 years of age and had continued musical training through the time of data collection (11-23 years of training; mean(SD)= 16.30(2.52) years), were included in Study 2. 10 non-musicians (mean(SD) = 25.8(4.1) years of age; 6 female), all of whom had less than 2 years of total musical training which could not have occurred either before 7 years of age or within the last 5 years (0-2 years of musical training; mean(SD)= 0.3(0.67) years of formal training) served as a comparison group. See (Boebinger et al., 2021) for additional details on this study.

##### fMRI task

The fMRI task used a mini-block design in which 2-second natural sound clips were repeated three times in a row (each mini-block for a given natural sound was 10.2 seconds in duration to allow for a sparse scanning sequence; see Data acquisition below). There were 192 natural sounds, including a set of 165 sounds frequently heard and recognizable from everyday life (originally from (S. Norman-Haignere et al., 2015)) as well as 27 additional music and drumming sound clips from different musical cultures (Boebinger et al., 2021). The sounds were categorized by independent raters via an online experiment into 14 different sound categories (conditions): English Speech, Foreign Speech, Human Vocalization, Human Non-Vocalization, Animal Vocalization, Animal Non-Vocalization, Nature, Mechanical, Song, Music, Environmental Sound, Drum, Foreign Music, Foreign Song. For the purposes of the present study, we compared responses to a subset of conditions, including three conditions that correspond to the three main conditions in Study 1 – language (English speech), foreign speech, and music (Western instrumental music) – as well as the following additional conditions: foreign music (non-Western instrumental music), song (Western vocal music, in English), foreign song (non-Western vocal music, not in English), and drums. The foreign music clips came from non-Western musical cultures that varied in tonality and rhythmic complexity (e.g., Indian raga, Balinese gamelan, Chinese opera, Mongolian throat singing, Jewish klezmer, Ugandan lamellophone music). Stimuli were ramped on and off with a 25-ms linear ramp. Either the second or third repetition of each sound was 12 dB quieter (67 dB SPL, compared to the 75 dB SPL for all other sounds), and participants were instructed to press an in-scanner button box when they heard the quieter sound. Participants completed 48 runs across three scanning sessions (16 per session). Each run consisted of 24 mini-blocks and five silent rest periods (also 10.2 seconds in duration), which were distributed evenly throughout each run, for a total run duration of 5.5 min. Each of the 192 stimuli was presented in two mini-blocks per scanning session, for a total of six mini-block repetitions per stimulus over the three scanning sessions. Stimulus order was randomly permuted across runs and across participants. Participants wore MR-compatible earphones in the scanner and were provided with an MR-compatible button box to respond to task prompts.

### fMRI data acquisition, preprocessing, and modeling

For ease of comparison with other published studies using these data, we chose to retain the preprocessing and modeling pipeline used previously. Furthermore, because each study included its own control group (adults with typical amounts of music experience), we could compare each study’s results to previous studies with typical adults (e.g., (Chen et al., 2023)). Replicating previous findings for these control groups is also useful to demonstrate the robustness of those findings to preprocessing and analysis choices (cf. (Botvinik-Nezer et al., 2020)). Finally, it is worth noting that in several past studies, we have established that the activation maps for the language localizer obtained through different pipelines (e.g., an SPM-based pipeline vs. a Freesurfer-based pipeline; (Wolna et al., 2026)) are extremely similar. Because all the preprocessing and modeling choices in the pipelines used here are standard, we do not expect the results to differ if an alternative pipeline were used.

#### Study 1 (Children)

##### Data acquisition

MRI data were collected at the Athinoula A. Martinos Imaging Center of the McGovern Institute for Brain Research at MIT on a 3T Siemens scanner using one of two custom 32-channel head coils made for children (Keil et al., 2011), a 12-channel coil, or the standard Siemens 32-channel head coil. Whole-head, high-resolution T1-weighted structural images were collected for each participant (voxel size = 1.33 mm (12ch coil) or 1 mm (all others) isotropic, 128 (12-ch coil) or 176 (all others) interleaved sagittal slices, FoV = 256 mm (adult coil) or 192 mm (pediatric coils); GRAPPA parallel imaging, acceleration factor of 3). Whole-brain (minus cerebellum) functional BOLD data were collected in 30 (12-ch coil) or 32 (all others) near-axial slices (aligned with the anterior/posterior commissure) in an interleaved order (TR = 2.0 s, TE = 30 ms, flip angle = 90°, FoV = 192 mm (pediatric coils) or 256 mm (standard 32-ch coil), voxel size = 3×3×4mm (12-ch coil) or 3 mm isotropic (all others)). PACE was used to adjust the position of the gradients based on the participant’s motion one TR back (Thesen et al., 2000).

##### Preprocessing

fMRI data were preprocessed and analyzed using SPM12 (release 7487), CONN EvLab module (release 19b) and custom MATLAB scripts. Each participant’s functional and structural data were converted from DICOM to NIfTI format. All functional scans were co-registered and resampled using B-spline interpolation to the first scan of the first session. Potential outlier scans were identified from the resulting subject-motion estimates as well as from BOLD signal indicators using default thresholds in the CONN pre-processing pipeline (5 st. dev. or more above the mean in global BOLD signal change or framewise displacement values above 0.9 mm). Functional and structural data were independently normalized into a common space (the Montreal Neurological Institute (MNI) template, IXI549Space) using the SPM12 unified segmentation and normalization procedure with a reference functional image computed as the mean functional image after realignment across all time points, omitting outlier scans. The output data were resampled to a common bounding box between MNI-space coordinates (−90, −126, and −72) and (90, 90, and 108), using 2 mm isotropic voxels and fourth-order spline interpolation for the functional data and 1 mm isotropic voxels and tri-linear interpolation for the structural data. Lastly, the functional data were smoothed spatially using spatial convolution with a 4 mm full-width half-maximum (FWHM) Gaussian kernel.

##### First-level modeling

Effects were estimated using a general linear model (GLM) in which each experimental condition (language, foreign speech, and music) and the response periods were modeled with a boxcar function convolved with the canonical hemodynamic response function (HRF) (fixation was modeled implicitly). Temporal autocorrelations in the BOLD signal timeseries were accounted for by a combination of high-pass filtering with a 128 s cutoff and whitening using an AR (0.2) model (first-order autoregressive model linearized around the coefficient a = 0.2) to approximate the observed covariance of the functional data in the context of restricted maximum likelihood (ReML) estimation. In addition to main condition effects, other model parameters in the GLM design included first-order temporal derivatives for each condition (for modeling spatial variability in the HRF delays) as well as nuisance regressors to control for the effect on the BOLD signal of slow linear drifts, subject-motion parameters, and outlier scans.

#### Study 2 (Musicians)

##### Data acquisition

MRI data were collected at the Athinoula A. Martinos Imaging Center of the McGovern Institute for Brain Research at MIT on a 3T Siemens Prisma with a 32-channel head coil. Whole-head, high-resolution T1-weighted structural images were collected for each participant (TR = 2.53 s, voxel size = 1 mm isotropic, 176 slices, 256 × 256 matrix). Whole-brain functional BOLD data were collected in 48 roughly axial slices (3 mm thick; oriented parallel to the anterior-posterior commissure line) having an in-plane resolution of 2.1 × 2.1mm (96 × 96 matrix, 0.3-mm slice gap). A simultaneous multislice (SMS) acceleration factor of 4 was used to minimize acquisition time (TA = 1.02 s). Each volume acquisition lasted 1 s, and the 2-s stimuli were presented during periods of silence between each acquisition, with a 200-ms buffer of silence before and after stimulus presentation, so one brain volume was collected every 3.4 s (1 s + 2 s + 0.2*2 s; TR = 3.4 s, TA = 1.02 s, TE = 33 ms, flip angle = 90°, 4 discarded initial acquisitions).

##### Preprocessing

Preprocessing and data analysis were performed using FSL software and custom Matlab scripts. Functional volumes were motion-corrected, slice-time-corrected, skull-stripped, linearly detrended, and aligned to each participant’s anatomical image (using FLIRT and BBRegister; (Greve & Fischl, 2009; Jenkinson & Smith, 2001)). Motion correction and function-to-anatomical registration were done separately for each run. Preprocessed data were then resampled to the cortical surface reconstruction computed by FreeSurfer (Dale et al., 1999) and smoothed on the surface using a 3-mm full-width half-maximum (FWHM) kernel to improve signal-to-noise ratio (SNR). The data were then downsampled to a 2-mm isotropic grid on the FreeSurfer-flattened cortical surface.

##### First-level modeling

The response to each of the 14 sound categories was estimated using GLMdenoise (Kay et al., 2013). Each condition was modeled as a boxcar regressor convolved with a hemodynamic response function (HRF), the shape of which was estimated from the data. To remove correlated noise across voxels, GLMdenoise also included a data-driven set of noise regressors derived from principal components of a noise-pool timeseries, with the optimal number of components selected via cross-validation. Beta weights were estimated by bootstrapping across runs, with the median across bootstrap samples taken as the final parameter estimate

### Functional regions of interest

In both studies, for each participant, we defined their language regions so we could probe the responses in these regions to the critical music condition(s). To do so, we used the contrast between language and foreign speech, which was present in both studies (Study 1: [mental, social, physical] > [foreign]; Study 2: [English] > [foreign]). This contrast (and similar contrasts) between a language condition and a perceptually similar condition where little/no linguistic content is discernible (such as the foreign speech condition here, or backwards speech / sequences of nonwords in other studies) has been extensively validated in past work and shown to reliably identify language-selective areas ((Fedorenko et al., 2010; Malik-Moraleda et al., 2022; H. Olson et al., 2023; Scott et al., 2017) *inter alia;* see (Fedorenko et al., 2024) for a review). Recent work has shown that this approach works well in children as young as 3-4 years old (Hiersche et al., 2024; Ozernov-Palchik et al., 2026).

To define these language functional regions of interest (fROIs), we used the group-constrained subject-specific approach (Fedorenko et al., 2010), where a set of masks (parcels) is derived from a group-level representation of the data for the same contrast and then combined with individual activation maps; within each parcel, subject-specific voxels are selected using a portion of the data (Study 1: leave-one-run-out validation; Study 2: 6-fold cross-validation with fROIs defined on 5/6 of runs, responses measured on the held-out 1/6). Here, we used five parcels derived from language localizer data in 220 adult participants (independent of the adult samples in the current study) and used in much past work (e.g., (Fedorenko et al., 2020; Ivanova et al., 2020; Jouravlev et al., 2020; Mollica et al., 2020; H. Olson et al., 2023; Ozernov-Palchik et al., 2026; Paunov et al., 2019; Pritchett et al., 2018; Shain et al., 2023). These included three regions in the left frontal cortex (falling within the inferior frontal gyrus (IFG) and its orbital part (IFGorb), and within the middle frontal gyrus (MFG)) and two regions on the lateral surface of the temporal cortex (AntTemp and (PostTemp). Individual fROIs were defined by selecting the 10% of voxels within each parcel with the highest t-value for the language contrast. Response magnitude (Study 1: percent BOLD signal change; Study 2: beta weights) were then estimated and averaged across the voxels in each fROI for the conditions of interest using held-out data (not used to define the fROIs). For completeness, we also defined language fROIs in the right hemisphere homotopic parcels (using the left-hemisphere parcels mirrored onto the right hemisphere).

### Statistical analyses

Statistical analyses were conducted in R (R version 4.2.1 (2022-06-23); R Core Team, 2022), using the average activation (response magnitude) per condition per fROI per participant. Conditions were compared using linear mixed effects models (fit with maximum likelihood) predicting response magnitude. To test for network-level main effects of condition and age, we used: lmer(magnitude ~ condition + age + (1|ROI) + (1|participant), REML = FALSE), where conditions are language, foreign speech, and music (as well as foreign music, song, foreign song, and drum for some analyses in Study 2), age is in years (included in the model with children only), and ROI (region of interest within the network) and participant are included as random effects. Significance was determined at a level of p < 0.05. We additionally tested for an interaction between condition and age in analyses with the children only (magnitude ~ condition*age + (1|ROI) + (1|participant)). To test for an effect of experience in both studies (Study 1: Children vs. Adults; Study 2: Musicians vs. Non-musicians), we included a condition * group interaction term (magnitude ~ condition*group + (1|ROI) + (1|participant)). Individual regions were also examined using the same models without ROI modeled as a random effect; significance was determined at a level of p < 0.05 Bonferroni corrected for the number of ROIs (five core left-hemisphere language regions, and five right-hemisphere language regions in the supplementary analyses; p < 0.01). Omnibus Type III tests were used for group x effect interactions to test for main effects. Estimated marginal means were adjusted for age, as applicable. Pairwise comparisons were Tukey-corrected.

## RESULTS

### Language areas show minimal responses to music in children

To determine whether the language network responds to music in childhood, we compared the brain’s responses to three conditions (language, foreign speech, and music) in 4-16-year-old children (n=80) with a group of adults (n=24).

We first examined the adults’ responses to these conditions, expecting to find higher responses to language than music or foreign speech in individually-defined language regions, as demonstrated previously (Chen et al., 2023; Deen et al., 2015; Fedorenko et al., 2011). Indeed, consistent with prior work, condition was a significant predictor of response magnitude across the left-hemisphere language network in adults (**Figure 1A; Table S1**): relative to language, responses were lower for foreign speech (Est=-0.851, SE=0.073, t(331.79)= −11.57, p<0.001) and music (Est=−1.045, SE=0.073, t(331.79)= −14.22, p<0.001). Pairwise comparisons of estimated marginal means showed that all condition differences were significant: language > foreign speech (Δ=0.851, p<0.001), language > music (Δ=1.045, p<0.001), and foreign speech > music (Δ=0.195, p=0.024) (**Figure S3A**). Responses to language were also higher than responses to foreign speech and music in each of the five left-hemisphere language regions individually (**Figure 1C**; **Table S2**) and similar patterns were observed in right-hemisphere homotopes of language areas (**Figure S1**; **Tables S4-S5**).

**Figure 1.**
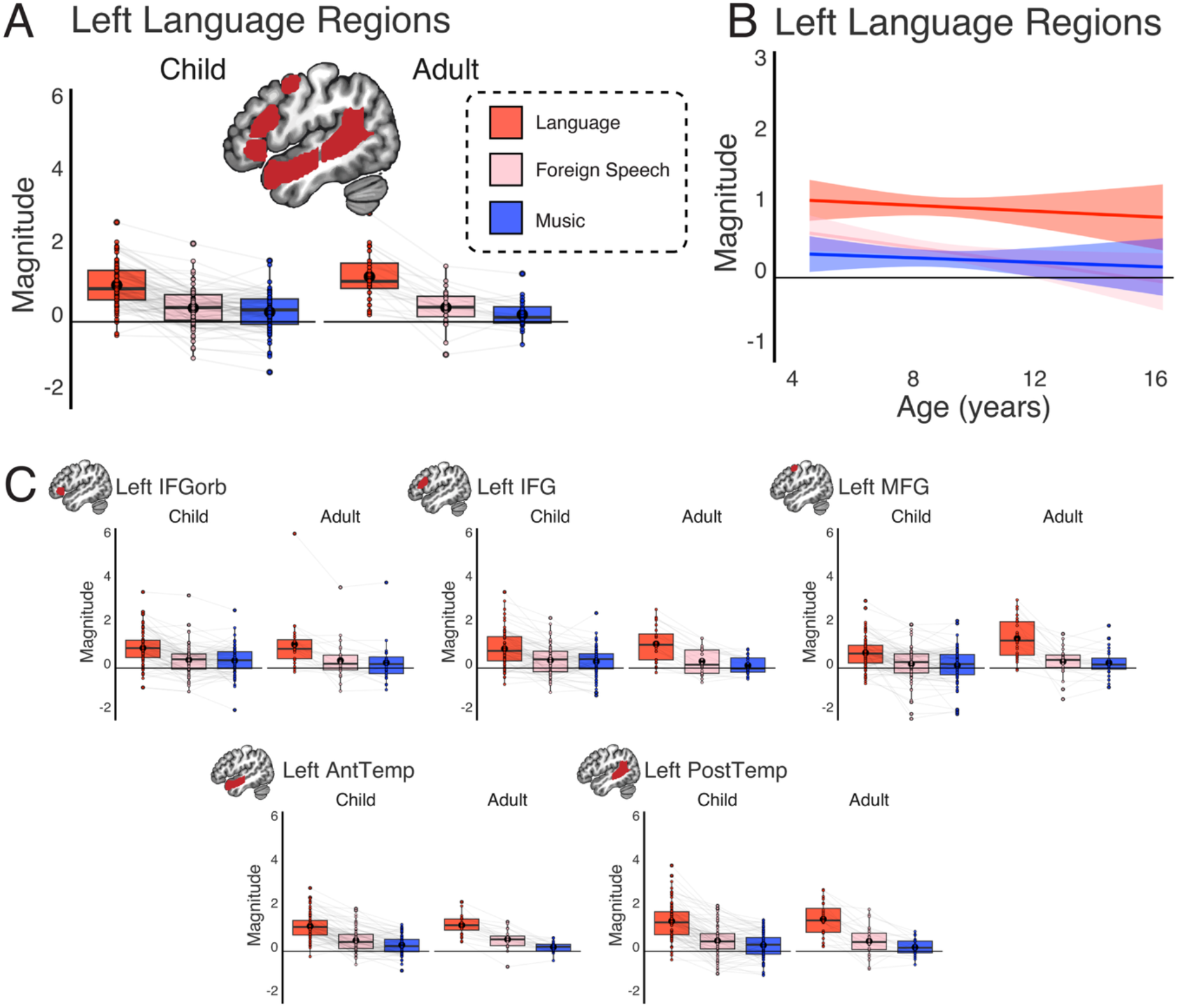
Children’s responses to language and music in canonical language regions, relative to adult controls. **(A)** Boxplots show response magnitude for language (red), foreign speech (pink) and music (blue), in children (n=80) and adults (n=24), averaged across five individually-defined left-hemisphere language regions. Dots show individual participants’ responses; light gray lines connect the same participant; large black dots show means. Language parcels (search spaces) shown on the brain in red; participants’ fROIs for language were defined as the top 10% of voxels responding to language>foreign speech in each parcel (responses per condition were extracted using independent data; Methods). **(B)** Trendlines show response magnitude per condition across age (years) in children (n=80), averaged across the five individually-defined left-hemisphere language regions. **(C)** Boxplots show response magnitude per condition for each language region.

Critically, the same pattern was observed in children (**Figure 1A**). Responses were lower for both foreign speech (Est=-0.626, SE=0.036, t(1115.93)=-17.23, p<0.001; this contrast reproduces the results reported in Ozernov-Palchik, O’Brien et al., 2026) and music (Est=-0.738, SE=0.036, t(1115.93)= −20.300, p<0.001) relative to the language condition in canonical left-hemisphere language regions. Age was not a significant predictor in the model. Pairwise comparisons of estimated marginal means (adjusted for age) showed that all condition differences were significant: language > foreign speech (Δ=0.626, p<0.001), language > music (Δ=0.738, p<0.001), and foreign speech > music (Δ=0.112, p=0.006) (**Figure S3A**). As in adults, responses to language were also higher than responses to foreign speech and music in each of the five left-hemisphere language regions (**Figure 1C; Table S2**).

When both children and adults were included in the same model, there was a main effect of condition (F(2, 1451.92)= 299.81; p<0.001)) and a significant interaction between condition and age group (F(2, 1451.92)= 8.33; p<0.001)). Follow-up estimated marginal means analyses showed that this interaction was driven by numerically higher responses to the language condition in adults than children (Δ=0.236, p=0.051), which is consistent with prior evidence of increasing response magnitude to language across childhood (Hiersche et al., 2024), including from a study that included the sample studied here (Ozernov-Palchik et al., 2026). Critically, however, there was no evidence of a difference between children and adults in response magnitude for the foreign speech or music conditions (**Table S3**). Thus, in children – as in adults – language regions’ responses to music were low and smaller than the responses to incomprehensible foreign speech, demonstrating a lack of responsivity to music.

### No developmental change in language areas’ responses to music across childhood

We next tested for a developmental change in the language regions’ responses to music across childhood. We fit another linear mixed-effects model predicting response magnitude in children’s left-hemisphere language regions, this time testing for an interaction between condition and age. In addition to a main effect of condition (F(2, 1115.93)=14.20, p<0.001), there was a significant interaction between condition and age (F(2, 1115.93)=4.229, p=0.015). Posthoc simple-slopes analyses revealed that age was negatively associated with response magnitude in the foreign speech condition (Est=-0.059, SE=0.026, t(108)= −2.278, p=0.025), but not for language (Est=-0.021, SE=0.026, t(108)=-0.797, p=0.427) or music (Est=-0.016, SE=0.026, t(108)=-0.604, p=0.547). Tukey-adjusted slope contrasts showed that while the slope was more negative for foreign speech than language and music, there was no evidence for a difference in slope between language and music (**Figure 1B**; see **Table S1** for full model results). By 4 years of age, the language network already shows very low responses to music which do not strengthen nor weaken over childhood; in other words, there is no evidence of developmental change across childhood in the language network’s selectivity for language over music.

### Extensive music experience does not affect language areas’ selectivity for language over music

Study 1 showed that the language network’s response to music was low and similar across children and adults, who differ in their amount of experience with music. We next compared responses to language, foreign speech, and music in the language network of adult musicians who have had continuous music training since childhood (n=10) and adults with no or minimal music training (n=10).

As expected, the non-musician group exhibited lower responses to foreign speech (Est= −0.223, SE= 0.023, t(135.73)= −9.703, p<0.001) and music (Est= −0.297, SE= 0.023, t(135.73)= −12.905, p<0.001) than language, and pairwise comparisons of estimated marginal means showed that all condition differences were significant: language > foreign speech (Δ=0.223, p<0.001), language > music (Δ=0.297, p<0.001), and foreign speech > music (Δ=0.074, p= 0.005). Critically, in musicians, we observed the same pattern. Condition was a significant predictor of response magnitude across the left-hemisphere language network (**Figure 2A**): relative to language, responses were lower for foreign speech (Est= −0.160, SE= 0.028, t(135.82)= −5.638, p<0.001) and music (Est= −0.239, SE= 0.028, t(135.82)= −8.422, p<0.001). Pairwise comparisons of estimated marginal means showed that all condition differences were significant: language > foreign speech (Δ=0.160, p<0.001), language > music (Δ=0.239, p<0.001), and foreign speech > music (Δ=0.079, p=0.018). Responses to language were also higher than responses to foreign speech and music in each of the five left-hemisphere language regions for both musicians and non-musicians (in left IFGorb and MFG regions in musicians, this effect was not significant for language > foreign speech, but the direction of the effect was the same; **Figure 2B**; **Table S7**). When both musicians and non-musicians were included in the same model, there was a main effect of condition (F(2, 275.84)= 112.36; p<0.001), but no effect of group nor an interaction between group and condition (**Table S6**). Thus, in highly-trained musicians, similar to adults with typical levels of musical experience, language regions do not respond to music. Similar patterns were observed in right-hemisphere homotopes of language areas (**Figure S2**; **Tables S10-S11**).

**Figure 2.**
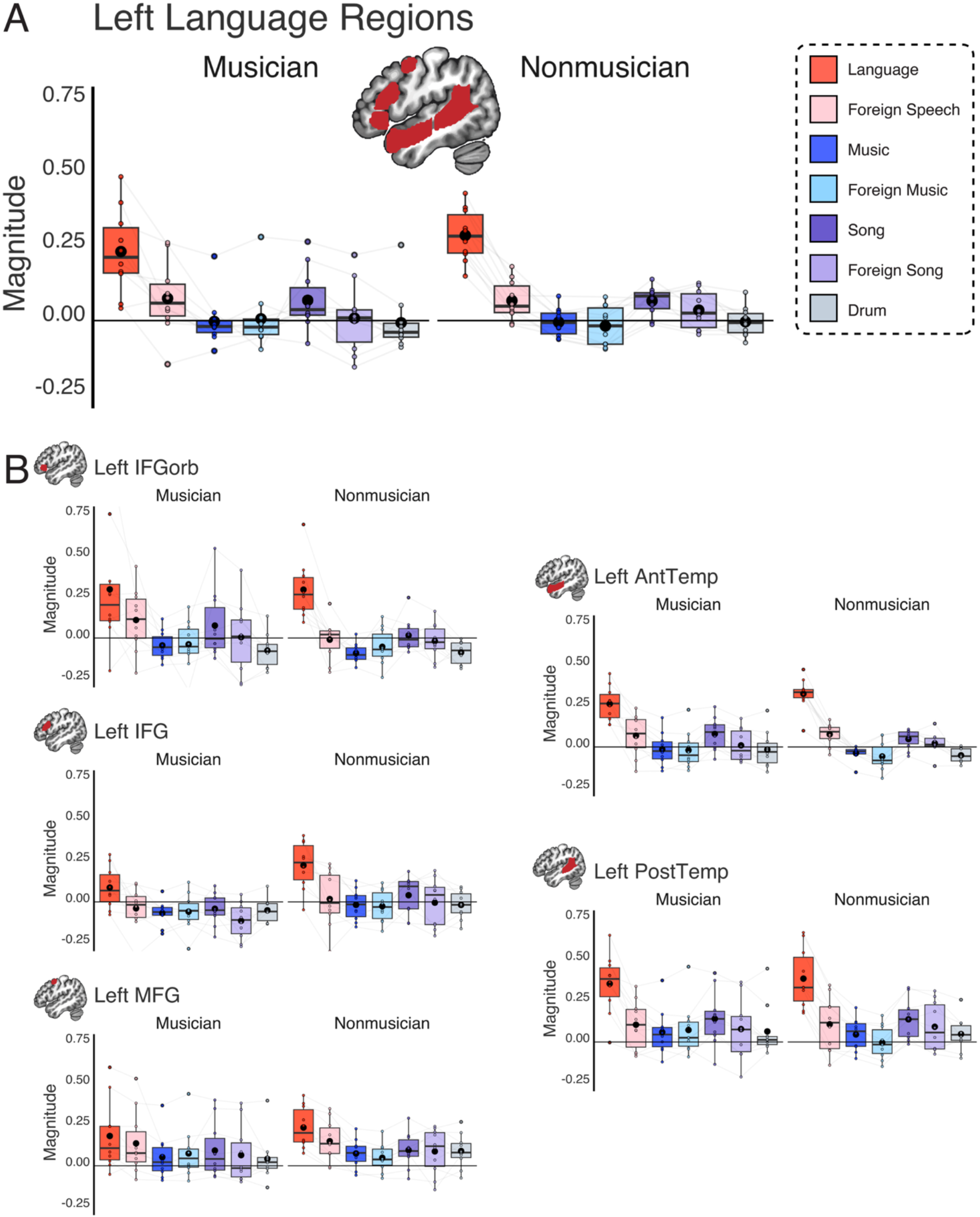
Musicians’ responses to language and music in canonical language regions, relative to adult non-musicians. **(A)** Boxplots show response magnitude for language (red), foreign speech (pink), music (blue), foreign music (light blue), songs (purple), foreign songs (light purple), and drumming (gray), in musicians (n=10) and non-musicians (n=10), averaged across five individually-defined left-hemisphere language regions. Dots show individual participants’ responses; light gray lines connect the same participant; large black dots show means. Language parcels (search spaces) shown on the brain in red; participants’ fROIs for language were defined as the top 10% of voxels responding to language>foreign speech in each parcel (responses per condition were extracted using independent data; Methods). **(B)** Boxplots show response magnitude per condition for each language region.

In this study, we also had the opportunity to examine four additional music conditions: non-Western instrumental music (foreign music), vocal music in English (song), non-Western vocal music (foreign song), and drumming. The inclusion of the foreign instrumental and song conditions allowed us to test whether the language system was sensitive to the familiarity of music, particularly in the musicians with extensive training, and the drumming condition has minimal melodic features, thereby isolating the rhythmic features of music, which some have argued may share machinery with language (Alagöz et al., 2025; Fiveash et al., 2021; Heard & Lee, 2020). Past studies that have included these conditions have found that the music-selective areas respond strongly to all of these music conditions (S. Norman-Haignere et al., 2015), including in the same sample included here (Boebinger et al., 2021). Past work has also shown that the language system of adults with typical amounts of musical experience responds relatively strongly to an English song condition (as expected given that it contains comprehensible language), higher than responses to instrumental music or drumming (Chen et al., 2023).

With all seven conditions included in the model, there was again a main effect of condition (F(6, 675.9)= 71.32; p<0.001), but no effect of group nor an interaction between group and condition (**Table S8**). Pairwise comparisons of estimated marginal means revealed that language responses were higher than all other conditions, in both musicians and non-musicians (**Figure S3B; Table S9**). In addition, both musicians and non-musicians showed higher responses to foreign speech than music and drumming, and higher responses to songs than drumming. Non-musicians additionally showed significant differences for foreign speech > foreign music and song > music (musicians showed a numerical difference in the same direction), as well as song > foreign music. The higher responses to the song condition are unsurprising given that the lyrics were comprehensible to participants, though these responses were still lower than the language condition, in line with the findings reported in Chen et al. (2023). Together, these results show that language regions show low responses to music regardless of familiarity with that particular style of music, even in highly trained musicians.

## DISCUSSION

In adults with typical amounts of music experience, listening to music does not elicit a response in the language areas of the brain (Chen et al., 2023; Fedorenko et al., 2011; Rogalsky et al., 2011; Sueoka et al., 2024). Here we showed that this selectivity for language over music holds regardless of the amount of music experience. Language areas were strongly selective for language over music in children, who have substantially less experience with both language and music than a typical adult, and in highly trained musicians, who have far more music experience. In both groups, responses to music were low overall and in children, this selectivity was stable between 4 and 16 years of age, showing no evidence for developmental change. Together, these results argue against music experience shaping the language network’s selectivity for language over music.

### Developmental trajectory of language network selectivity

These results add to the growing body of evidence that, by the time children are fluently using language, their language network is selective for language relative to non-linguistic inputs and tasks. In particular, prior work has established that by early childhood, the language network already shows adult-like topography, including a strong left-hemispheric bias (Ozernov-Palchik et al., 2026), and dissociates from other networks that support cognitive functions like working memory (Hiersche et al., 2024) or theory of mind (Hiersche et al., 2026). Our results extend these findings to music – a high-level perceptual domain that shares critical early experiences with language.

Although this work found selectivity for language over music by 4 years of age, it leaves open the possibility that music may overlap with language processing earlier in life. Unlike the music areas, which show evidence of selectivity for music as early as one month of age (Kosakowski et al., 2023), the language areas cannot be selective for language relative to a condition like foreign or backwards speech until children begin mapping words to meaning (indeed, the infants in the same study did not show selectivity for speech; (Kosakowski et al., 2023); see also (Cristia et al., 2014)). The wordform-to-meaning mapping begins in infancy (Bergelson & Swingley, 2012), but it is not until toddlerhood that children acquire a vocabulary of a few hundred words and start combining them to infer and create complex meanings – an ability that continues to develop throughout childhood (Bloom, 1991; Bloom & Lahey, 1978; Frank et al., 2021). Indeed, though toddlers show the emergence of a topographically similar language system to adults, the magnitude of the language response is small (H. A. Olson et al., 2026) and its selectivity for language is not yet known. It is therefore possible that some of the similarities between language and music in early development – including exposure to both in utero (Birnholz & Benacerraf, 1983; Chorna et al., 2019; Graven & Browne, 2008), a domain-general capacity for auditory statistical learning (Saffran, 2003; Saffran et al., 1996), and the roles of language and music in infant care and bonding (Mehr et al., 2021; Savage et al., 2021) – would lead to shared neural underpinnings in early development. However, the present results suggest that language selectivity is robustly present by childhood.

### Music expertise and language network selectivity

Although language regions do not *typically* respond to music in mature or developing brains, music experience may still affect the selectivity of the language network. In particular, extensive experience with structured non-linguistic auditory inputs could lead to progressively more cortical tissue needed to support the relevant ability, and this expansion could encroach on the language system given the proximity of its temporal components to the auditory cortex.

Experience-dependent brain plasticity has been observed across domains, both in typical development (e.g., the specialization of the visual word form area for reading; (Baker et al., 2007; Dehaene & Cohen, 2011; McCandliss et al., 2003)) and in atypical development (e.g., responses in the visual cortex of congenitally blind individuals to higher-order cognition; (Saccone et al., 2024)). The language network also exhibits experience-dependent plasticity: responses to language continue to increase in magnitude across childhood, likely as a function of additional language experience (Hiersche et al., 2024; Ozernov-Palchik et al., 2026). Learning a new language leads to responsiveness to that language, even in adulthood (Malik-Moraleda et al., 2024; Wolna et al., 2024) and for hand-constructed languages (Malik-Moraleda et al., 2025). And although most natural languages rely on the auditory modality, the language network is flexible enough to accommodate languages that use other modalities (e.g., for sign or tactile languages; (Emmorey & McCullough, 2009; MacSweeney et al., 2008)). However, this experience-dependent plasticity has limits: for example, learning a new skill like computer coding does not recruit the language mechanisms (Ivanova et al., 2020; Liu et al., 2020), despite deep similarities between natural and programming languages (Fedorenko et al., 2019).

Here, we found that despite its similarities to language, music does not engage the language regions even in the case of early and extensive music training. These results generalized across Western music (the type of music that our musician population was most exposed to), non-Western music, and drumming. Interestingly, as reported in Boebinger et al. (2021), responses to these conditions are also similar between musicians and non-musicians in the music-selective areas, with both groups showing similarly high responses. Together, these results suggest that if extensive music training leads to changes in how the brain responds to music, these effects are located neither in the auditory music-selective areas nor in the language network. Finally, in both musicians and non-musicians, the language areas responded more strongly to language than to song, in line with prior work (Chen et al., 2023). This effect may be due to decreased attention to the linguistic content in songs, whose melodic and rhythmic structure may draw attentional resources away from the lyrics.

These results not only provide further evidence for the distinction between language and music processing in the brain, but also shed light on the limitations of the language system’s sensitivity to experience: even extensive experience with hierarchically structured auditory stimuli does not change the selectivity of the language areas. These findings align with prior clinical evidence from some musicians who experienced strokes but continued their music careers after developing severe aphasia (Basso & Capitani, 1985; Luria et al., 1965).

### Why doesn’t music engage language-processing mechanisms?

Language shares similarities with many non-linguistic domains: from math, logic, and computer code, to various social-communicative abilities, to music. Past work has demonstrated that between-domain similarities need not imply shared machinery (for review, see (Fedorenko et al., 2024)). This study provides additional evidence that a key feature of music – its hierarchical structure, which is also a signature of natural language – does not result in overlapping brain mechanisms, and establishes that this separation holds in childhood and persists despite extensive music training.

These results suggest that some critical differences between music and language push them toward distinct neural substrates. Indeed, language and music serve different functions: the core function of language is communicating propositional meanings (Gibson et al., 2019; Pinker & Bloom, 1990), whereas music is highly limited in the kinds of meanings it can express (e.g., (Jackendoff, 2009)). Early linguistic theorizing has emphasized the abstract nature of syntax (e.g., (Chomsky, 1957)) and has likely led to a slew of hypotheses about language sharing machinery with other domains that rely on hierarchical structures (e.g., (Carruthers, 2002; Fitch, 2014; Koechlin & Jubault, 2006; Patel, 2003)). However, empirical evidence now favors usage-based accounts of language (e.g., (Bybee, 2010; Goldberg, 1995)), where syntactic structures are deeply connected to and shaped by the meanings they express, and much of our language knowledge consists of idiosyncratic phrases and constructions. This optimization of language, including its grammar, for efficiently sharing meanings (Gibson et al., 2019) makes it fundamentally different from music and apparently different enough to draw on distinct machinery in the brain. The fact that other meaning-rich domains like math and logic do not engage the language-processing mechanisms (Ivanova et al., 2020; Kean et al., 2026; Liu et al., 2020; Monti et al., 2007, 2009) further suggests that it may be particular *kinds* of meanings that the language system is optimized for computing (e.g., meanings that have to do with the internal and external worlds; cf. more abstract relational meanings that mathematical or logic expressions carry; see, e.g., discussion in (Malik-Moraleda et al., 2025) and (Kean et al., 2026)). The function(s) of music have always been a mystery (Darwin, 1871; McDermott, 2008; Pinker, 1994) and remain debated (e.g., (Mehr et al., 2021; Savage et al., 2021)). However, thinking about what music is for in our species is likely critical for understanding not only the computations carried out by the music-selective areas, but also the relationship between these areas and various other cognitive and neural systems.

### Future directions

One important open question is whether areas that are selective for language by early childhood may respond to music earlier in development. Longitudinal neuroimaging with infants and toddlers can examine this possibility. Second, unlike Study 1, the stimuli in Study 2 consisted of short (2-second) clips: longer, more complex stimuli may offer a more sensitive test for detecting differences between musicians and non-musicians. Finally, generalizing these findings beyond the W.E.I.R.D. groups tested here remains an important goal (Henrich et al., 2010).

## Conclusion

Humans have a specialized system in the brain for processing language (Fedorenko et al., 2024). Despite numerous similarities between language and music, we found that this language system is not engaged in music perception either in childhood or in highly trained musicians. Thus, experience does not appear to affect the selectivity of the language system for language over music – instead, fundamental differences in the functions that language and music serve likely require distinct neural substrates to support these capacities.

## AUTHOR CONTRIBUTIONS

HAO: Designed research; Analyzed data; Wrote the paper. XJC: Analyzed data; Edited the paper. DB: Designed research; Performed research; Analyzed data; Edited the paper. HR: Designed research; Performed research; Edited the paper. EF: Designed research; Edited the paper.

## ACKNOWLEDGEMENTS

We would like to acknowledge the Athinoula A. Martinos Imaging Center at the McGovern Institute for Brain Research at MIT and its support team (Steve Shannon and Atsushi Takahashi), Rebecca Saxe, Nancy Kanwisher, and Josh McDermott for their contributions to the design of the original studies and supervising work on those studies, Elizabeth Lee for her help in preprocessing and analyzing the data from Study 1 in an updated pipeline, and the participants and families for making this research possible.

