## Supplement for "The language network does not support music perception across a range of music experience"

**Supplementary Materials**

#### Supplementary Figures

##### Figure S1: Responses in right language homotopic regions (Study 1)

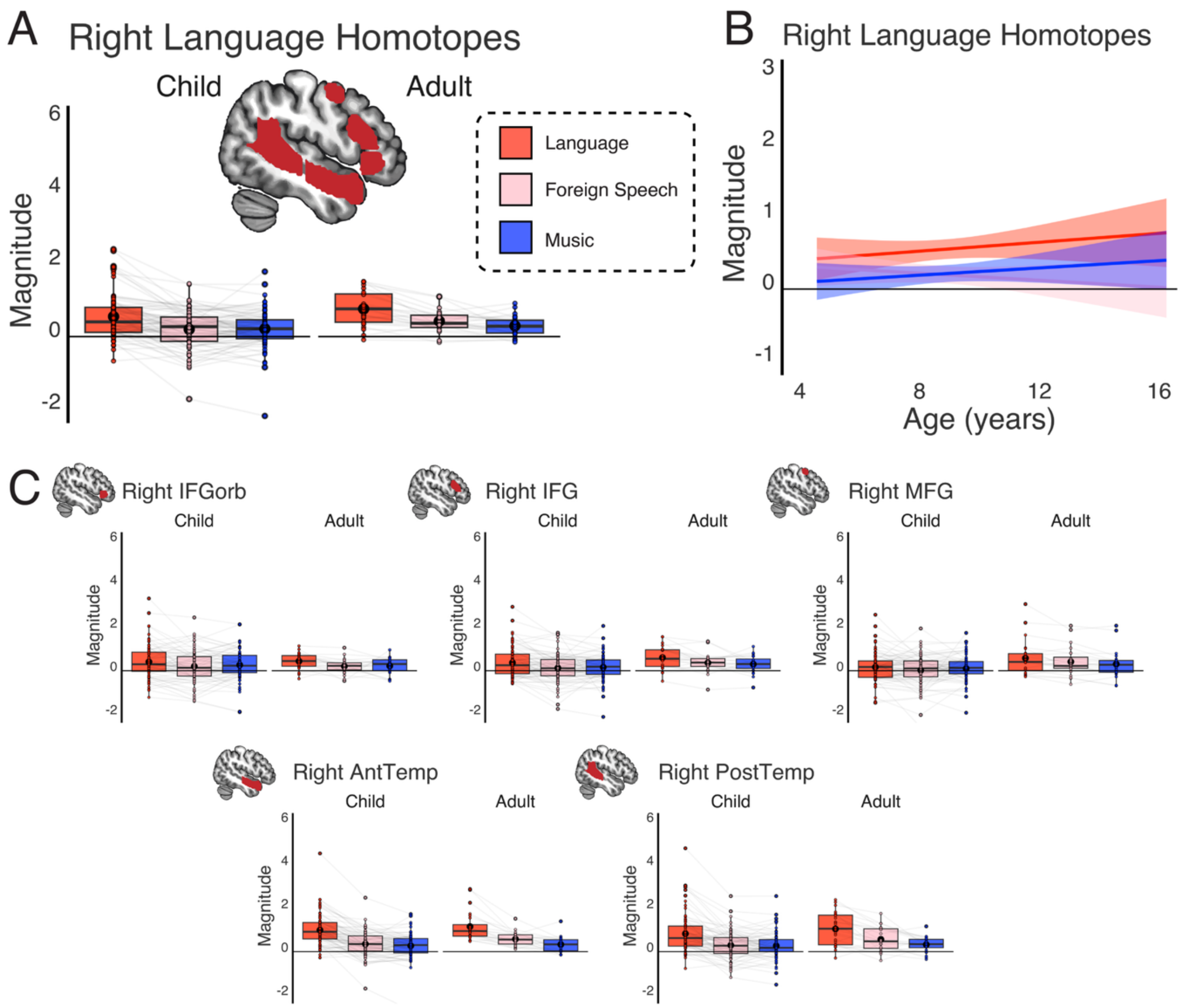

**Children's responses to language and music in right-hemisphere homotopes of language regions**. **(A)** Boxplots show response magnitude for language (red), foreign speech (pink) and music (blue), in children (n=80) and adults (n=24), averaged across five right-hemisphere homotopes of language regions. Dots show individual participants' responses; light gray lines connect the same participant; large black dots show means. Search spaces shown on the brain in red; participants' fROIs were defined as the top 10% of voxels responding to language>foreign speech in each search space (responses per condition were extracted from independent data). **(B)** Trendlines show response magnitude per condition across age (years) in children (N=80). **(C)** Boxplots show response magnitude per condition for each region.

**Figure S2: Responses in right language homotopic regions (Study 2)**

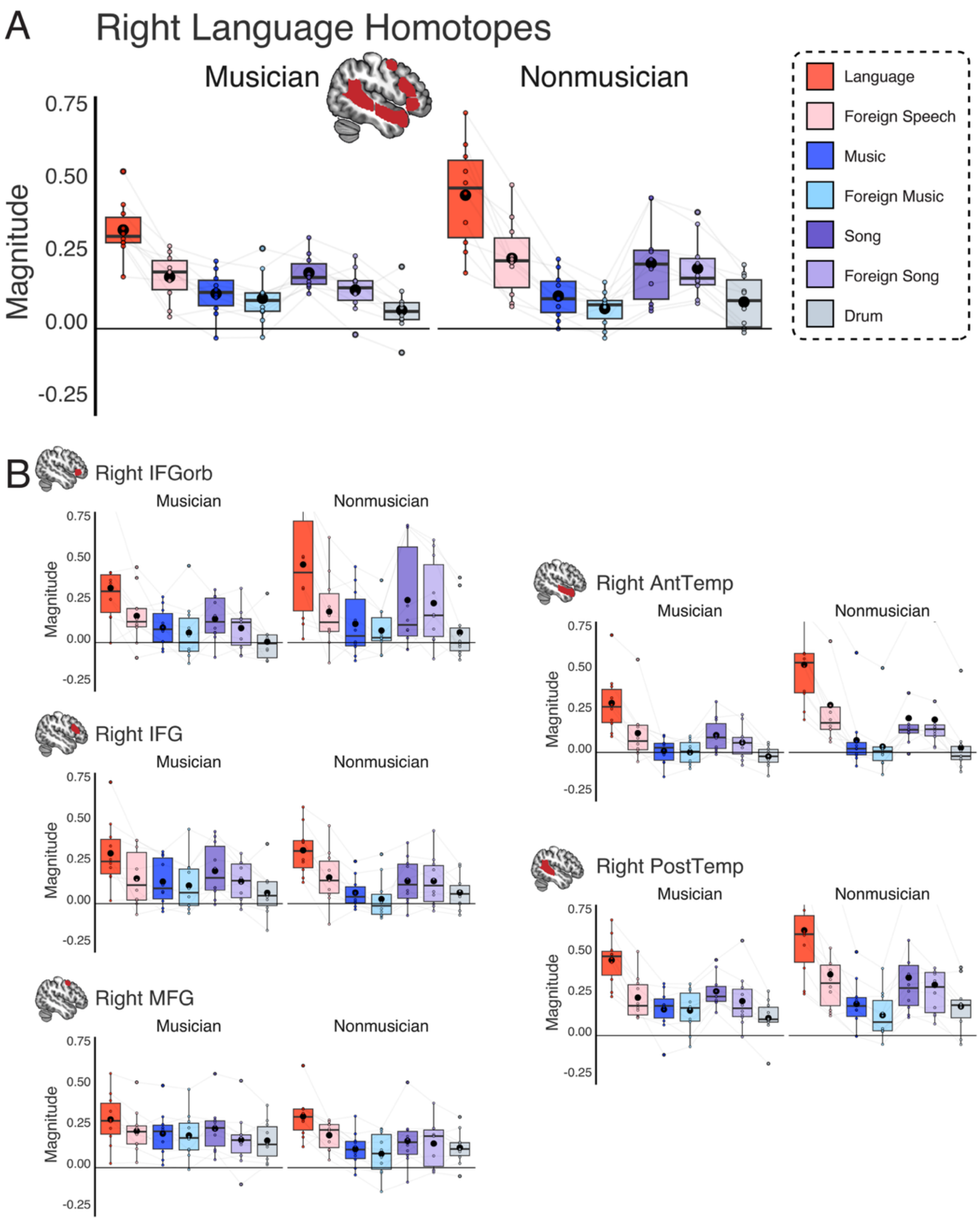

**Musician's responses to language and music in right-hemisphere homotopes of language regions**. **(A)** Boxplots show response magnitude for language (red), foreign speech (pink), music (blue), foreign music (light blue), songs (purple), foreign songs (light purple), and drumming (gray), in musicians (n=10) and non-musicians (n=10), averaged across the five right-hemisphere homotopes of language regions. Dots show individual participants' responses; light gray lines connect the same participant; large black dots show means. Search spaces shown on the brain in red; participants' fROIs were defined as the top 10% of voxels responding to language>foreign speech in each search space (responses per condition were extracted from independent data). **(B)** Boxplots show response magnitude per condition for each region.

**Figure S3: Heatmap of pairwise condition differences within each group**

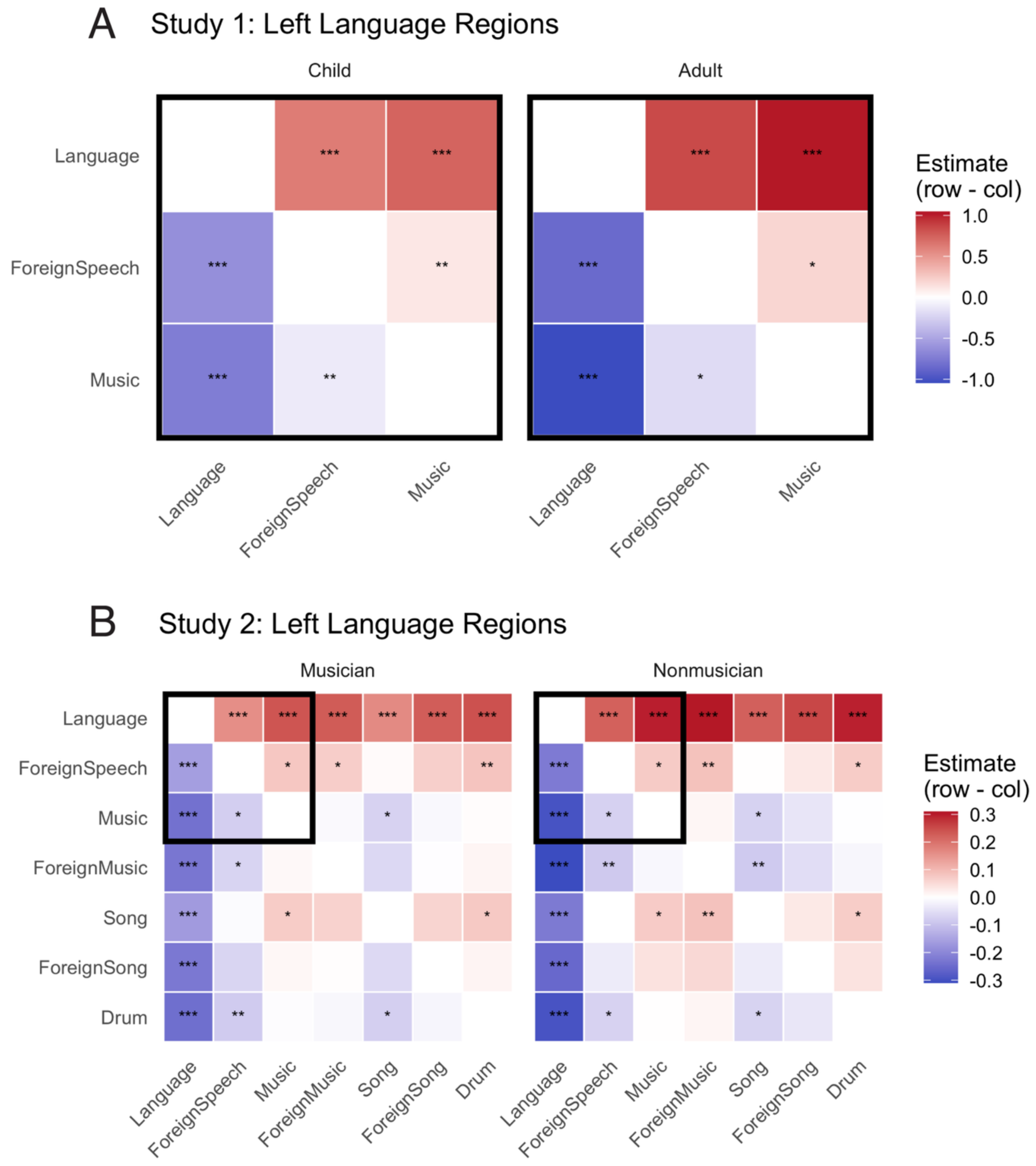

**Condition differences by group**. Heatmaps show pairwise comparisons of estimated marginal means for each group, in **(A)** Study 1 and **(B)** Study 2, for the mixed-effects models for the left-hemisphere language network including both groups (Magnitude~Condition*Group+(1|ROI)+(1|Subject)). Color indicates the condition difference for the row – column (red=positive; blue=negative). Stars indicate significance using Tukey-adjusted p-values (*p<0.05, **p<0.01, ***p<0.001). Black border highlights analogous conditions between Study 1 and Study 2.

### Supplementary Tables

#### Table S1: Mixed-effects model results for left-hemisphere language network (Study 1)

| Term | Estimate | SE | df | t | p | CI low | CI high |
|---|---|---|---|---|---|---|---|
| **CHILDREN** | Magnitude~Condition+Age+(1\|ROI)+(1\|Subject) | | | | | | |
| **Intercept** | 1.285 | 0.226 | 84.197 | 5.680 | **<0.001** | 0.835 | 1.735 |
| **Foreign Speech vs Language** | −0.626 | 0.036 | 1115.929 | −17.228 | **<0.001** | −0.698 | −0.555 |
| **Music vs Language** | −0.738 | 0.036 | 1115.929 | −20.300 | **<0.001** | −0.809 | −0.667 |
| **Age (years)** | −0.032 | 0.024 | 79.679 | −1.329 | 0.1876 | −0.079 | 0.016 |
| **ADULTS** | Magnitude~Condition+(1\|ROI)+(1\|Subject) | | | | | | |
| **Intercept** | 1.241 | 0.104 | 33.139 | 11.990 | **<0.001** | 1.031 | 1.452 |
| **Foreign Speech vs Language** | −0.851 | 0.073 | 331.788 | −11.572 | **<0.001** | −0.995 | −0.706 |
| **Music vs Language** | −1.045 | 0.073 | 331.788 | −14.220 | **<0.001** | −1.190 | −0.901 |
| **ADULTS VS CHILDREN** | Magnitude~Condition*Group+(1\|ROI)+(1\|Subject) | | | | | | |
| **Intercept** | 1.005 | 0.075 | 21.995 | 13.344 | **<0.001** | 0.849 | 1.161 |
| **Foreign Speech vs Language** | -0.626 | 0.037 | 1451.920 | -16.741 | **<0.001** | -0.700 | -0.553 |
| **Music vs Language** | -0.738 | 0.037 | 1451.912 | -19.726 | **<0.001** | -0.811 | -0.664 |
| **Adults vs Children** | 0.236 | 0.119 | 140.727 | 1.983 | **0.049** | 0.0008 | 0.472 |
| **Foreign Speech vs Language : Adults vs Children** | -0.224 | 0.078 | 1451.912 | -2.881 | **0.004** | -0.377 | -0.072 |
| **Music vs Language : Adults vs Children** | -0.307 | 0.078 | 1451.912 | -3.946 | **<0.001** | -0.460 | -0.155 |
| **CHILDREN** | Magnitude~Condition*Age+(1\|ROI)+(1\|Subject) | | | | | | |
| **Intercept** | 1.187 | 0.241 | 108.276 | 4.924 | **<0.001** | 0.709 | 1.665 |
| **Foreign Speech vs Language** | -0.288 | 0.148 | 1115.929 | -1.939 | 0.053 | -0.579 | 0.003 |
| **Music vs Language** | -0.782 | 0.148 | 1115.929 | -5.268 | **<0.001** | -1.073 | -0.491 |
| **Age (years)** | -0.021 | 0.026 | 106.175 | -0.803 | 0.424 | -0.072 | 0.030 |
| **Foreign Speech vs Language : Age** | -0.038 | 0.016 | 1115.929 | -2.352 | **0.019** | -0.070 | -0.006 |
| **Music vs Language : Age** | 0.005 | 0.016 | 1115.929 | 0.305 | 0.760 | -0.027 | 0.037 |

Significant p-values in bold ($p < 0.05$).

**Table S2: Mixed-effects model results for individual language regions (Study 1)**

| Term | Estimate | SE | df | t | p | CI low | CI high |
|---|---|---|---|---|---|---|---|
| **LH IFGorb** | | | | | | | |
| **CHILDREN** | Magnitude~Condition+Age+(1\|Subject) | | | | | | |
| **Intercept** | 1.434 | 0.279 | 82.979 | 5.145 | **<0.001** | 0.880 | 1.988 |
| **Foreign Speech vs Language** | −0.541 | 0.065 | 160.000 | −8.313 | **<0.001** | −0.670 | −0.413 |
| **Music vs Language** | −0.576 | 0.065 | 160.000 | −8.848 | **<0.001** | −0.705 | −0.448 |
| **Age (years)** | −0.059 | 0.030 | 80.000 | −1.939 | 0.0560 | −0.119 | 0.002 |
| **ADULTS** | Magnitude~Condition+(1\|Subject) | | | | | | |
| **Intercept** | 1.083 | 0.208 | 32.335 | 5.203 | **<0.001** | 0.659 | 1.507 |
| **Foreign Speech vs Language** | −0.740 | 0.137 | 48.000 | −5.396 | **<0.001** | −1.016 | −0.464 |
| **Music vs Language** | −0.840 | 0.137 | 48.000 | −6.128 | **<0.001** | −1.116 | −0.565 |
| **LH IFG** | | | | | | | |
| **CHILDREN** | Magnitude~Condition+Age+(1\|Subject) | | | | | | |
| **Intercept** | 1.263 | 0.279 | 83.219 | 4.530 | **<0.001** | 0.709 | 1.818 |
| **Foreign Speech vs Language** | −0.529 | 0.068 | 160.000 | −7.814 | **<0.001** | −0.662 | −0.395 |
| **Music vs Language** | −0.581 | 0.068 | 160.000 | −8.583 | **<0.001** | −0.715 | −0.447 |
| **Age (years)** | −0.043 | 0.030 | 80.000 | −1.422 | 0.1589 | −0.104 | 0.017 |
| **ADULTS** | Magnitude~Condition+(1\|Subject) | | | | | | |
| **Intercept** | 1.101 | 0.125 | 37.811 | 8.800 | **<0.001** | 0.848 | 1.355 |
| **Foreign Speech vs Language** | −0.794 | 0.101 | 48.000 | −7.843 | **<0.001** | −0.998 | −0.591 |
| **Music vs Language** | −0.984 | 0.101 | 48.000 | −9.713 | **<0.001** | −1.187 | −0.780 |
| **LH MFG** | | | | | | | |
| **CHILDREN** | Magnitude~Condition+Age+(1\|Subject) | | | | | | |
| **Intercept** | 0.958 | 0.297 | 83.714 | 3.225 | **0.0018** | 0.367 | 1.550 |
| **Foreign Speech vs Language** | −0.514 | 0.077 | 160.000 | −6.649 | **<0.001** | −0.667 | −0.361 |
| **Music vs Language** | −0.574 | 0.077 | 160.000 | −7.418 | **<0.001** | −0.726 | −0.421 |
| **Age (years)** | −0.030 | 0.032 | 80.000 | −0.920 | 0.3601 | −0.094 | 0.035 |
| **ADULTS** | Magnitude~Condition+(1\|Subject) | | | | | | |

| | | | | | | | |
|---|---|---|---|---|---|---|---|
| **Intercept** | 1.355 | 0.151 | 47.679 | 8.967 | **<0.001** | 1.051 | 1.659 |
| **Foreign Speech vs Language** | −1.062 | 0.150 | 48.000 | −7.063 | **<0.001** | −1.364 | −0.760 |
| **Music vs Language** | −1.118 | 0.150 | 48.000 | −7.438 | **<0.001** | −1.421 | −0.816 |
| **LH AntTemp** | | | | | | | |
| **CHILDREN** | Magnitude~Condition+Age+(1\|Subject) | | | | | | |
| **Intercept** | 0.978 | 0.205 | 83.247 | 4.769 | **<0.001** | 0.570 | 1.386 |
| **Foreign Speech vs Language** | −0.648 | 0.050 | 160.000 | −12.964 | **<0.001** | −0.747 | −0.549 |
| **Music vs Language** | −0.860 | 0.050 | 160.000 | −17.200 | **<0.001** | −0.958 | −0.761 |
| **Age (years)** | 0.019 | 0.022 | 80.000 | 0.871 | 0.3861 | −0.025 | 0.064 |
| **ADULTS** | Magnitude~Condition+(1\|Subject) | | | | | | |
| **Intercept** | 1.195 | 0.078 | 51.779 | 15.361 | **<0.001** | 1.039 | 1.351 |
| **Foreign Speech vs Language** | −0.649 | 0.082 | 48.000 | −7.895 | **<0.001** | −0.814 | −0.484 |
| **Music vs Language** | −0.997 | 0.082 | 48.000 | −12.124 | **<0.001** | −1.162 | −0.831 |
| **LH PostTemp** | | | | | | | |
| **CHILDREN** | Magnitude~Condition+Age+(1\|Subject) | | | | | | |
| **Intercept** | 1.792 | 0.277 | 83.591 | 6.459 | **<0.001** | 1.240 | 2.343 |
| **Foreign Speech vs Language** | −0.899 | 0.071 | 160.000 | −12.658 | **<0.001** | −1.039 | −0.758 |
| **Music vs Language** | −1.099 | 0.071 | 160.000 | −15.478 | **<0.001** | −1.239 | −0.959 |
| **Age (years)** | −0.046 | 0.030 | 80.000 | −1.541 | 0.1273 | −0.106 | 0.014 |
| **ADULTS** | Magnitude~Condition+(1\|Subject) | | | | | | |
| **Intercept** | 1.471 | 0.127 | 51.439 | 11.578 | **<0.001** | 1.216 | 1.726 |
| **Foreign Speech vs Language** | −1.007 | 0.134 | 48.000 | −7.537 | **<0.001** | −1.276 | −0.738 |
| **Music vs Language** | −1.286 | 0.134 | 48.000 | −9.627 | **<0.001** | −1.555 | −1.018 |

Significant p-values in bold, Bonferroni corrected for the number of ROIs ($p < 0.01$).

#### Table S3: Pairwise contrasts: Group differences within each effect (Study 1)

| Term | Estimate | SE | df | CI low | CI high | T ratio | p |
|---|---|---|---|---|---|---|---|
| | | | | LH Language Network | | | |
| CHILDREN - ADULTS | Magnitude~Condition*Group+(1\|ROI)+(1\|Subject) | | | | | | |
| Language | -0.236 | 0.120 | 142.435 | -0.473 | 0.0007 | -1.971 | 0.051 |
| Foreign Speech | -0.012 | 0.120 | 142.435 | -0.249 | 0.2259 | -0.100 | 0.921 |
| Music | 0.071 | 0.120 | 142.435 | -0.166 | 0.308 | 0.592 | 0.555 |

##### Table S4: Mixed-effects model results for right-hemisphere homotopic language areas (Study 1)

| Term | Estimate | SE | df | t | p | CI low | CI high |
|---|---|---|---|---|---|---|---|
| **CHILDREN** | Magnitude~Condition+Age+(1\|ROI)+(1\|Subject) | | | | | | |
| **Intercept** | 0.461 | 0.235 | 82.060 | 1.957 | 0.0537 | −0.008 | 0.929 |
| **Foreign Speech vs Language** | −0.354 | 0.036 | 1,115.945 | −9.777 | **<0.001** | −0.425 | −0.283 |
| **Music vs Language** | −0.340 | 0.036 | 1,115.945 | −9.383 | **<0.001** | −0.411 | −0.269 |
| **Age (years)** | 0.011 | 0.025 | 79.586 | 0.434 | 0.6654 | −0.038 | 0.059 |
| **ADULTS** | Magnitude~Condition+(1\|ROI)+(1\|Subject) | | | | | | |
| **Intercept** | 0.767 | 0.096 | 15.838 | 8.018 | **<0.001** | 0.564 | 0.970 |
| **Foreign Speech vs Language** | −0.340 | 0.058 | 331.852 | −5.811 | **<0.001** | −0.455 | −0.225 |
| **Music vs Language** | −0.469 | 0.058 | 331.852 | −8.023 | **<0.001** | −0.584 | −0.354 |
| **ADULTS VS CHILDREN** | Magnitude~Condition*Group+(1\|ROI)+(1\|Subject) | | | | | | |
| **Intercept** | 0.555 | 0.086 | 12.412 | 6.413 | **<0.001** | 0.367 | 0.742 |
| **Foreign Speech vs Language** | -0.354 | 0.035 | 1451.949 | -10.000 | **<0.001** | -0.424 | -0.285 |
| **Music vs Language** | -0.340 | 0.035 | 1451.949 | -9.598 | **<0.001** | -0.410 | -0.271 |
| **Adults vs Children** | 0.212 | 0.113 | 140.006 | 1.874 | 0.063 | -0.012 | 0.437 |
| **Foreign Speech vs Language : Adults vs Children** | 0.015 | 0.074 | 1451.949 | 0.197 | 0.844 | -0.130 | 0.159 |
| **Music vs Language : Adults vs Children** | -0.129 | 0.074 | 1451.949 | -1.750 | 0.080 | -0.274 | 0.016 |

Significant p-values in bold ($p < 0.05$).

#### Table S5: Mixed-effects model results for individual right-hemisphere homotopic language areas (Study 1)

| Term | Estimate | SE | df | t | p | CI low | CI high |
|---|---|---|---|---|---|---|---|
| **RH IFGorb** | | | | | | | |
| **CHILDREN** | Magnitude~Condition+Age+(1\|Subject) | | | | | | |
| **Intercept** | 0.423 | 0.304 | 82.305 | 1.389 | 0.1684 | −0.183 | 1.029 |
| **Foreign Speech vs Language** | −0.209 | 0.063 | 160.000 | −3.326 | **0.0011** | −0.332 | −0.085 |
| **Music vs Language** | −0.143 | 0.063 | 160.000 | −2.273 | 0.0243 | −0.266 | −0.019 |
| **Age** | −0.001 | 0.033 | 80.000 | −0.035 | 0.9724 | −0.067 | 0.065 |
| **ADULTS** | Magnitude~Condition+(1\|Subject) | | | | | | |
| **Intercept** | 0.444 | 0.073 | 62.687 | 6.099 | **<0.001** | 0.299 | 0.590 |
| **Foreign Speech vs Language** | −0.236 | 0.088 | 48.000 | −2.690 | **0.0098** | −0.413 | −0.060 |
| **Music vs Language** | −0.219 | 0.088 | 48.000 | −2.497 | 0.0160 | −0.396 | −0.043 |
| **RH IFG** | | | | | | | |
| **CHILDREN** | Magnitude~Condition+Age+(1\|Subject) | | | | | | |
| **Intercept** | 0.345 | 0.256 | 83.674 | 1.348 | 0.1812 | −0.164 | 0.853 |
| **Foreign Speech vs Language** | −0.244 | 0.066 | 160.000 | −3.684 | **<0.001** | −0.374 | −0.113 |
| **Music vs Language** | −0.210 | 0.066 | 160.000 | −3.180 | **0.0018** | −0.341 | −0.080 |
| **Age** | 0.003 | 0.028 | 80.000 | 0.120 | 0.9050 | −0.052 | 0.059 |
| **ADULTS** | Magnitude~Condition+(1\|Subject) | | | | | | |
| **Intercept** | 0.609 | 0.092 | 45.030 | 6.626 | **<0.001** | 0.424 | 0.795 |
| **Foreign Speech vs Language** | −0.239 | 0.088 | 48.000 | −2.728 | **0.0089** | −0.415 | −0.063 |
| **Music vs Language** | −0.305 | 0.088 | 48.000 | −3.485 | **0.0011** | −0.481 | −0.129 |
| **RH MFG** | | | | | | | |
| **CHILDREN** | Magnitude~Condition+Age+(1\|Subject) | | | | | | |
| **Intercept** | 0.244 | 0.265 | 83.948 | 0.920 | 0.3600 | −0.283 | 0.771 |
| **Foreign Speech vs Language** | −0.147 | 0.071 | 160.000 | −2.067 | 0.0403 | −0.287 | −0.007 |
| **Music vs Language** | −0.066 | 0.071 | 160.000 | −0.927 | 0.3555 | −0.206 | 0.074 |
| **Age** | −0.008 | 0.029 | 80.000 | −0.287 | 0.7745 | −0.066 | 0.049 |

<table>
<tr><td>ADULTS</td><td colspan="7">Magnitude~Condition+(1|Subject)</td></tr>
<tr><td>Intercept</td><td>0.582</td><td>0.140</td><td>28.106</td><td>4.146</td><td><0.001</td><td>0.294</td><td>0.869</td></tr>
<tr><td>Foreign Speech vs Language</td><td>−0.164</td><td>0.068</td><td>48.000</td><td>−2.420</td><td>0.0194</td><td>−0.300</td><td>−0.028</td></tr>
<tr><td>Music vs Language</td><td>−0.270</td><td>0.068</td><td>48.000</td><td>−3.986</td><td>0.0002</td><td>−0.406</td><td>−0.134</td></tr>
<tr><td colspan="8">RH AntTemp</td></tr>
<tr><td>CHILDREN</td><td colspan="7">Magnitude~Condition+Age+(1|Subject)</td></tr>
<tr><td>Intercept</td><td>0.616</td><td>0.249</td><td>83.185</td><td>2.479</td><td>0.0152</td><td>0.122</td><td>1.110</td></tr>
<tr><td>Foreign Speech vs Language</td><td>−0.641</td><td>0.060</td><td>160.000</td><td>−10.678</td><td><0.001</td><td>−0.759</td><td>−0.522</td></tr>
<tr><td>Music vs Language</td><td>−0.723</td><td>0.060</td><td>160.000</td><td>−12.059</td><td><0.001</td><td>−0.842</td><td>−0.605</td></tr>
<tr><td>Age</td><td>0.042</td><td>0.027</td><td>80.000</td><td>1.567</td><td>0.1210</td><td>−0.011</td><td>0.096</td></tr>
<tr><td>ADULTS</td><td colspan="7">Magnitude~Condition+(1|Subject)</td></tr>
<tr><td>Intercept</td><td>1.155</td><td>0.096</td><td>49.179</td><td>12.047</td><td><0.001</td><td>0.962</td><td>1.348</td></tr>
<tr><td>Foreign Speech vs Language</td><td>−0.577</td><td>0.098</td><td>48.000</td><td>−5.913</td><td><0.001</td><td>−0.773</td><td>−0.381</td></tr>
<tr><td>Music vs Language</td><td>−0.828</td><td>0.098</td><td>48.000</td><td>−8.479</td><td><0.001</td><td>−1.024</td><td>−0.631</td></tr>
<tr><td colspan="8">RH PostTemp</td></tr>
<tr><td>CHILDREN</td><td colspan="7">Magnitude~Condition+Age+(1|Subject)</td></tr>
<tr><td>Intercept</td><td>0.675</td><td>0.288</td><td>83.419</td><td>2.346</td><td>0.0213</td><td>0.103</td><td>1.248</td></tr>
<tr><td>Foreign Speech vs Language</td><td>−0.532</td><td>0.072</td><td>160.000</td><td>−7.394</td><td><0.001</td><td>−0.674</td><td>−0.390</td></tr>
<tr><td>Music vs Language</td><td>−0.558</td><td>0.072</td><td>160.000</td><td>−7.759</td><td><0.001</td><td>−0.700</td><td>−0.416</td></tr>
<tr><td>Age</td><td>0.017</td><td>0.031</td><td>80.000</td><td>0.541</td><td>0.5899</td><td>−0.045</td><td>0.079</td></tr>
<tr><td>ADULTS</td><td colspan="7">Magnitude~Condition+(1|Subject)</td></tr>
<tr><td>Intercept</td><td>1.045</td><td>0.116</td><td>44.651</td><td>8.999</td><td><0.001</td><td>0.811</td><td>1.279</td></tr>
<tr><td>Foreign Speech vs Language</td><td>−0.483</td><td>0.110</td><td>48.000</td><td>−4.400</td><td><0.001</td><td>−0.704</td><td>−0.262</td></tr>
<tr><td>Music vs Language</td><td>−0.724</td><td>0.110</td><td>48.000</td><td>−6.597</td><td><0.001</td><td>−0.945</td><td>−0.503</td></tr>
</table>

Significant p-values in bold, Bonferroni corrected for the number of ROIs ($p < 0.01$).

#### Table S6: Mixed-effects model results for left-hemisphere language network; 3 conditions (Study 2)

| Term | Estimate | SE | df | t | p | CI low | CI high |
|---|---|---|---|---|---|---|---|
| **MUSICIANS** | Magnitude~Condition+(1\|ROI)+(1\|Subject) | | | | | | |
| **Intercept** | 0.236 | 0.043 | 15.079 | 5.465 | **<0.001** | 0.144 | 0.328 |
| **Foreign Speech vs Language** | −0.160 | 0.028 | 135.824 | −5.638 | **<0.001** | −0.216 | −0.104 |
| **Music vs Language** | −0.239 | 0.028 | 135.824 | −8.422 | **<0.001** | −0.295 | −0.183 |
| **NON-MUSICIANS** | Magnitude~Condition+(1\|ROI)+(1\|Subject) | | | | | | |
| **Intercept** | 0.291 | 0.029 | 14.417 | 9.881 | **<0.001** | 0.228 | 0.355 |
| **Foreign Speech vs Language** | −0.223 | 0.023 | 135.727 | −9.703 | **<0.001** | −0.269 | −0.178 |
| **Music vs Language** | −0.297 | 0.023 | 135.727 | −12.905 | **<0.001** | −0.343 | −0.251 |
| **NON-MUSICIANS VS MUSICIANS** | Magnitude~Condition*Group+(1\|ROI)+(1\|Subject) | | | | | | |
| **Intercept** | 0.236 | 0.036 | 23.574 | 6.551 | **<0.001** | 0.162 | 0.311 |
| **Foreign Speech vs Language** | -0.160 | 0.026 | 275.843 | -6.144 | **<0.001** | -0.211 | -0.109 |
| **Music vs Language** | -0.239 | 0.0261 | 275.843 | -9.178 | **<0.001** | -0.290 | -0.188 |
| **Non-musicians vs Musicians** | 0.055 | 0.042 | 35.655 | 1.330 | 0.192 | -0.029 | 0.140 |
| **Foreign Speech vs Language : Non-musicians vs Musicians** | -0.063 | 0.0369 | 275.843 | -1.715 | 0.088 | -0.136 | 0.009 |
| **Music vs Language : Non-musicians vs Musicians** | -0.058 | 0.037 | 275.843 | -1.569 | 0.118 | -0.130 | 0.015 |

Significant p-values in bold ($p < 0.05$).

**Table S7: Mixed-effects model results for individual language regions; 3 conditions (Study 2)**

| Term | Estimate | SE | df | t | p | CI low | CI high |
|---|---|---|---|---|---|---|---|
| **LH IFGorb** | | | | | | | |
| **MUSICIANS** | Magnitude~Condition+(1\|Subject) | | | | | | |
| **Intercept** | 0.296 | 0.074 | 23.203 | 3.983 | **<0.001** | 0.142 | 0.449 |
| **Foreign Speech vs Language** | −0.187 | 0.083 | 20.000 | −2.260 | 0.0351 | −0.359 | −0.014 |
| **Music vs Language** | −0.341 | 0.083 | 20.000 | −4.127 | **<0.001** | −0.513 | −0.169 |
| **NON-MUSICIANS** | Magnitude~Condition+(1\|Subject) | | | | | | |
| **Intercept** | 0.294 | 0.038 | 25.053 | 7.718 | **<0.001** | 0.216 | 0.372 |
| **Foreign Speech vs Language** | −0.304 | 0.045 | 20.000 | −6.812 | **<0.001** | −0.397 | −0.211 |
| **Music vs Language** | −0.386 | 0.045 | 20.000 | −8.654 | **<0.001** | −0.479 | −0.293 |
| **LH IFG** | | | | | | | |
| **MUSICIANS** | Magnitude~Condition+(1\|Subject) | | | | | | |
| **Intercept** | 0.087 | 0.034 | 21.491 | 2.598 | **0.0166** | 0.017 | 0.157 |
| **Foreign Speech vs Language** | −0.126 | 0.035 | 20.000 | −3.574 | **0.0019** | −0.200 | −0.053 |
| **Music vs Language** | −0.155 | 0.035 | 20.000 | −4.399 | **<0.001** | −0.229 | −0.082 |
| **NON-MUSICIANS** | Magnitude~Condition+(1\|Subject) | | | | | | |
| **Intercept** | 0.222 | 0.042 | 22.202 | 5.314 | **<0.001** | 0.136 | 0.309 |
| **Foreign Speech vs Language** | −0.205 | 0.045 | 20.000 | −4.549 | **<0.001** | −0.299 | −0.111 |
| **Music vs Language** | −0.240 | 0.045 | 20.000 | −5.331 | **<0.001** | −0.334 | −0.146 |
| **LH MFG** | | | | | | | |
| **MUSICIANS** | Magnitude~Condition+(1\|Subject) | | | | | | |
| **Intercept** | 0.181 | 0.055 | 13.094 | 3.315 | **0.0055** | 0.063 | 0.299 |
| **Foreign Speech vs Language** | −0.045 | 0.034 | 20.000 | −1.311 | 0.2049 | −0.116 | 0.027 |
| **Music vs Language** | −0.129 | 0.034 | 20.000 | −3.761 | **0.0012** | −0.200 | −0.057 |
| **NON-MUSICIANS** | Magnitude~Condition+(1\|Subject) | | | | | | |
| **Intercept** | 0.233 | 0.034 | 13.817 | 6.855 | **<0.001** | 0.160 | 0.306 |
| **Foreign Speech vs Language** | −0.083 | 0.023 | 20.000 | −3.570 | **0.0019** | −0.132 | −0.035 |

| | | | | | | | |
|---|---|---|---|---|---|---|---|
| **Music vs Language** | −0.158 | 0.023 | 20.000 | −6.786 | **<0.001** | −0.206 | −0.109 |
| **LH AntTemp** | | | | | | | |
| **MUSICIANS** | Magnitude~Condition+(1\|Subject) | | | | | | |
| **Intercept** | 0.262 | 0.033 | 16.405 | 7.950 | **<0.001** | 0.192 | 0.332 |
| **Foreign Speech vs Language** | −0.193 | 0.028 | 20.000 | −6.952 | **<0.001** | −0.251 | −0.135 |
| **Music vs Language** | −0.276 | 0.028 | 20.000 | −9.928 | **<0.001** | −0.334 | −0.218 |
| **NON-MUSICIANS** | Magnitude~Condition+(1\|Subject) | | | | | | |
| **Intercept** | 0.323 | 0.022 | 23.091 | 14.955 | **<0.001** | 0.279 | 0.368 |
| **Foreign Speech vs Language** | −0.246 | 0.024 | 20.000 | −10.295 | **<0.001** | −0.296 | −0.196 |
| **Music vs Language** | −0.361 | 0.024 | 20.000 | −15.080 | **<0.001** | −0.411 | −0.311 |
| **LH PostTemp** | | | | | | | |
| **MUSICIANS** | Magnitude~Condition+(1\|Subject) | | | | | | |
| **Intercept** | 0.354 | 0.047 | 14.807 | 7.536 | **<0.001** | 0.254 | 0.454 |
| **Foreign Speech vs Language** | −0.249 | 0.035 | 20.000 | −7.045 | **<0.001** | −0.323 | −0.176 |
| **Music vs Language** | −0.295 | 0.035 | 20.000 | −8.326 | **<0.001** | −0.369 | −0.221 |
| **NON-MUSICIANS** | Magnitude~Condition+(1\|Subject) | | | | | | |
| **Intercept** | 0.385 | 0.046 | 12.994 | 8.415 | **<0.001** | 0.286 | 0.484 |
| **Foreign Speech vs Language** | −0.278 | 0.028 | 20.000 | −9.827 | **<0.001** | −0.337 | −0.219 |
| **Music vs Language** | −0.340 | 0.028 | 20.000 | −12.004 | **<0.001** | −0.399 | −0.281 |

Significant p-values in bold, Bonferroni corrected for the number of ROIs ($p < 0.01$).

**Table S8: Mixed-effects model results for left-hemisphere language network; 7 conditions (Study 2)**

| Term | Estimate | SE | df | t | p | CI low | CI high |
|---|---|---|---|---|---|---|---|
| **MUSICIANS** | Magnitude~Condition+(1\|ROI)+(1\|Subject) | | | | | | |
| **Intercept** | 0.236 | 0.042 | 16.943 | 5.625 | **<0.001** | 0.147 | 0.325 |
| **Foreign Speech vs Language** | −0.160 | 0.025 | 335.941 | −6.472 | **<0.001** | −0.209 | −0.111 |
| **Music vs Language** | −0.239 | 0.025 | 335.941 | −9.668 | **<0.001** | −0.288 | −0.191 |
| **Foreign Music vs Language** | −0.230 | 0.025 | 335.941 | −9.300 | **<0.001** | −0.279 | −0.181 |
| **Song vs Language** | −0.167 | 0.025 | 335.941 | −6.737 | **<0.001** | −0.215 | −0.118 |
| **Foreign Song vs Language** | −0.228 | 0.025 | 335.941 | −9.200 | **<0.001** | −0.276 | −0.179 |
| **Drum vs Language** | −0.244 | 0.025 | 335.941 | −9.844 | **<0.001** | −0.292 | −0.195 |
| **NON-MUSICIANS** | Magnitude~Condition+(1\|ROI)+(1\|Subject) | | | | | | |
| **Intercept** | 0.291 | 0.029 | 15.441 | 10.200 | **<0.001** | 0.231 | 0.352 |
| **Foreign Speech vs Language** | −0.223 | 0.020 | 335.906 | −10.997 | **<0.001** | −0.263 | −0.183 |
| **Music vs Language** | −0.297 | 0.020 | 335.906 | −14.626 | **<0.001** | −0.337 | −0.257 |
| **Foreign Music vs Language** | −0.310 | 0.020 | 335.906 | −15.266 | **<0.001** | −0.350 | −0.270 |
| **Song vs Language** | −0.224 | 0.020 | 335.906 | −11.036 | **<0.001** | −0.264 | −0.184 |
| **Foreign Song vs Language** | −0.256 | 0.020 | 335.906 | −12.592 | **<0.001** | −0.296 | −0.216 |
| **Drum vs Language** | −0.295 | 0.020 | 335.906 | −14.545 | **<0.001** | −0.335 | −0.255 |
| **NON-MUSICIANS VS MUSICIANS** | Magnitude~Condition*Group+(1\|ROI)+(1\|Subject) | | | | | | |
| **Intercept** | 0.236 | 0.035 | 24.129 | 6.775 | **<0.001** | 0.164 | 0.308 |
| **Foreign Speech vs Language** | -0.160 | 0.023 | 675.947 | -6.999 | **<0.001** | -0.205 | -0.115 |
| **Music vs Language** | -0.239 | 0.023 | 675.947 | -10.455 | **<0.001** | -0.284 | -0.194 |
| **Foreign Music vs Language** | -0.230 | 0.023 | 675.947 | -10.057 | **<0.001** | -0.275 | -0.185 |
| **Song vs Language** | -0.167 | 0.023 | 675.947 | -7.284 | **<0.001** | -0.212 | -0.122 |
| **Foreign Song vs Language** | -0.228 | 0.023 | 675.947 | -9.948 | **<0.001** | -0.273 | -0.183 |
| **Drum vs Language** | -0.244 | 0.023 | 675.947 | -10.644 | **<0.001** | -0.288 | -0.199 |
| **Non-musicians vs Musicians** | 0.055 | 0.039 | 39.016 | 1.416 | 0.165 | -0.024 | 0.135 |
| **Foreign Speech vs Language : Non-musicians vs Musicians** | -0.063 | 0.032 | 675.947 | -1.953 | 0.051 | -0.127 | 0.000 |

| | | | | | | | |
|---|---|---|---|---|---|---|---|
| **Music vs Language : Non-musicians vs Musicians** | -0.058 | 0.032 | 675.947 | -1.787 | 0.074 | -0.121 | 0.006 |
| **Foreign Music vs Language : Non-musicians vs Musicians** | -0.080 | 0.032 | 675.947 | -2.470 | **0.014** | -0.143 | -0.016 |
| **Song vs Language : Non-musicians vs Musicians** | -0.057 | 0.032 | 675.947 | -1.775 | 0.076 | -0.121 | 0.006 |
| **Foreign Song vs Language : Non-musicians vs Musicians** | -0.028 | 0.032 | 675.947 | -0.868 | 0.386 | -0.092 | 0.035 |
| **Drum vs Language : Non-musicians vs Musicians** | -0.052 | 0.032 | 675.947 | -1.602 | 0.110 | -0.115 | 0.012 |

Significant p-values in bold ($p < 0.05$).

#### Table S9: Pairwise contrasts: Group differences within each effect; 7 conditions (Study 2)

| Term | Estimate | SE | df | CI low | CI high | T ratio | p |
|---|---|---|---|---|---|---|---|
| LH Language Network | | | | | | | |
| MUSICIAN - NONMUSICIAN | Magnitude~Condition*Group+(1\|ROI)+(1\|Subject) | | | | | | |
| Language | -0.055 | 0.040 | 41.513 | -0.137 | 0.026 | -1.376 | 0.176 |
| Foreign Speech | 0.008 | 0.040 | 41.513 | -0.074 | 0.089 | 0.193 | 0.848 |
| Music | 0.002 | 0.040 | 41.513 | -0.079 | 0.084 | 0.059 | 0.953 |
| Foreign Music | 0.024 | 0.040 | 41.513 | -0.057 | 0.106 | 0.608 | 0.547 |
| Song | 0.002 | 0.040 | 41.513 | -0.079 | 0.083 | 0.050 | 0.960 |
| Foreign Song | -0.027 | 0.040 | 41.513 | -0.109 | 0.054 | -0.679 | 0.501 |
| Drum | -0.004 | 0.040 | 41.513 | -0.085 | 0.078 | -0.089 | 0.929 |

**Table S10: Mixed-effects model results for right-hemisphere homotopic language areas; 3 conditions (Study 2)**

| Term | Estimate | SE | df | t | p | CI low | CI high |
|---|---|---|---|---|---|---|---|
| **MUSICIANS** | Magnitude~Condition+(1\|ROI)+(1\|Subject) | | | | | | |
| Intercept | 0.340 | 0.033 | 15.857 | 10.403 | **<0.001** | 0.271 | 0.409 |
| Foreign Speech vs Language | −0.162 | 0.030 | 135.581 | −5.454 | **<0.001** | −0.221 | −0.103 |
| Music vs Language | −0.219 | 0.030 | 135.581 | −7.391 | **<0.001** | −0.278 | −0.161 |
| **NON-MUSICIANS** | Magnitude~Condition+(1\|ROI)+(1\|Subject) | | | | | | |
| Intercept | 0.460 | 0.054 | 14.659 | 8.474 | **<0.001** | 0.344 | 0.575 |
| Foreign Speech vs Language | −0.217 | 0.037 | 135.810 | −5.915 | **<0.001** | −0.290 | −0.145 |
| Music vs Language | −0.347 | 0.037 | 135.810 | −9.448 | **<0.001** | −0.420 | −0.274 |
| **NON-MUSICIANS VS MUSICIANS** | Magnitude~Condition*Group+(1\|ROI)+(1\|Subject) | | | | | | |
| **Intercept** | 0.340 | 0.042 | 26.871 | 8.054 | **<0.001** | 0.253 | 0.427 |
| **Foreign Speech vs Language** | -0.162 | 0.034 | 275.786 | -4.757 | **<0.001** | -0.229 | -0.095 |
| **Music vs Language** | -0.219 | 0.034 | 275.786 | -6.446 | **<0.001** | -0.286 | -0.152 |
| **Non-musicians vs Musicians** | 0.120 | 0.051 | 39.887 | 2.364 | **0.023** | 0.017 | 0.222 |
| **Foreign Speech vs Language : Non-musicians vs Musicians** | -0.055 | 0.048 | 275.786 | -1.152 | 0.250 | -0.150 | 0.039 |
| **Music vs Language : Non-musicians vs Musicians** | -0.128 | 0.048 | 275.786 | -2.655 | **0.008** | -0.222 | -0.033 |

Significant p-values in bold ($p < 0.05$).

**Table S11: Mixed-effects model results for individual right-hemisphere homotopic language areas; 3 conditions (Study 2)**

| Term | Estimate | SE | df | t | p | CI low | CI high |
|---|---|---|---|---|---|---|---|
| **RH IFGorb** | | | | | | | |
| **MUSICIANS** | Magnitude~Condition+(1\|Subject) | | | | | | |
| **Intercept** | 0.334 | 0.055 | 18.680 | 6.079 | **<0.001** | 0.219 | 0.449 |
| **Foreign Speech vs Language** | −0.171 | 0.052 | 20.000 | −3.284 | **0.0037** | −0.280 | −0.062 |
| **Music vs Language** | −0.242 | 0.052 | 20.000 | −4.646 | **<0.001** | −0.351 | −0.133 |
| **NON-MUSICIANS** | Magnitude~Condition+(1\|Subject) | | | | | | |
| **Intercept** | 0.478 | 0.081 | 12.947 | 5.916 | **<0.001** | 0.303 | 0.652 |
| **Foreign Speech vs Language** | −0.288 | 0.050 | 20.000 | −5.817 | **<0.001** | −0.392 | −0.185 |
| **Music vs Language** | −0.363 | 0.050 | 20.000 | −7.324 | **<0.001** | −0.466 | −0.260 |
| **RH IFG** | | | | | | | |
| **MUSICIANS** | Magnitude~Condition+(1\|Subject) | | | | | | |
| **Intercept** | 0.307 | 0.052 | 14.418 | 5.906 | **<0.001** | 0.196 | 0.418 |
| **Foreign Speech vs Language** | −0.154 | 0.038 | 20.000 | −4.064 | **<0.001** | −0.233 | −0.075 |
| **Music vs Language** | −0.173 | 0.038 | 20.000 | −4.578 | **<0.001** | −0.252 | −0.094 |
| **NON-MUSICIANS** | Magnitude~Condition+(1\|Subject) | | | | | | |
| **Intercept** | 0.326 | 0.042 | 15.648 | 7.701 | **<0.001** | 0.236 | 0.415 |
| **Foreign Speech vs Language** | −0.166 | 0.034 | 20.000 | −4.879 | **<0.001** | −0.237 | −0.095 |
| **Music vs Language** | −0.259 | 0.034 | 20.000 | −7.626 | **<0.001** | −0.330 | −0.188 |
| **RH MFG** | | | | | | | |
| **MUSICIANS** | Magnitude~Condition+(1\|Subject) | | | | | | |
| **Intercept** | 0.295 | 0.044 | 12.968 | 6.734 | **<0.001** | 0.200 | 0.390 |
| **Foreign Speech vs Language** | −0.071 | 0.027 | 20.000 | −2.638 | 0.0158 | −0.127 | −0.015 |
| **Music vs Language** | −0.086 | 0.027 | 20.000 | −3.199 | **0.0045** | −0.143 | −0.030 |
| **NON-MUSICIANS** | Magnitude~Condition+(1\|Subject) | | | | | | |
| **Intercept** | 0.313 | 0.033 | 16.106 | 9.439 | **<0.001** | 0.243 | 0.384 |
| **Foreign Speech vs Language** | −0.115 | 0.028 | 20.000 | −4.189 | **<0.001** | −0.173 | −0.058 |

| | | | | | | | |
|---|---|---|---|---|---|---|---|
| **Music vs Language** | −0.200 | 0.028 | 20.000 | −7.275 | **<0.001** | −0.258 | −0.143 |
| **RH AntTemp** | | | | | | | |
| **MUSICIANS** | Magnitude~Condition+(1\|Subject) | | | | | | |
| **Intercept** | 0.302 | 0.046 | 18.535 | 6.539 | **<0.001** | 0.205 | 0.399 |
| **Foreign Speech vs Language** | −0.184 | 0.044 | 20.000 | −4.226 | **<0.001** | −0.275 | −0.093 |
| **Music vs Language** | −0.293 | 0.044 | 20.000 | −6.734 | **<0.001** | −0.384 | −0.202 |
| **NON-MUSICIANS** | Magnitude~Condition+(1\|Subject) | | | | | | |
| **Intercept** | 0.537 | 0.074 | 12.979 | 7.297 | **<0.001** | 0.378 | 0.696 |
| **Foreign Speech vs Language** | −0.247 | 0.045 | 20.000 | −5.443 | **<0.001** | −0.342 | −0.152 |
| **Music vs Language** | −0.462 | 0.045 | 20.000 | −10.185 | **<0.001** | −0.557 | −0.368 |
| **RH PostTemp** | | | | | | | |
| **MUSICIANS** | Magnitude~Condition+(1\|Subject) | | | | | | |
| **Intercept** | 0.462 | 0.041 | 18.037 | 11.340 | **<0.001** | 0.376 | 0.547 |
| **Foreign Speech vs Language** | −0.229 | 0.037 | 20.000 | −6.110 | **<0.001** | −0.307 | −0.151 |
| **Music vs Language** | −0.302 | 0.037 | 20.000 | −8.042 | **<0.001** | −0.380 | −0.223 |
| **NON-MUSICIANS** | Magnitude~Condition+(1\|Subject) | | | | | | |
| **Intercept** | 0.644 | 0.078 | 13.788 | 8.261 | **<0.001** | 0.476 | 0.811 |
| **Foreign Speech vs Language** | −0.270 | 0.053 | 20.000 | −5.070 | **<0.001** | −0.381 | −0.159 |
| **Music vs Language** | −0.451 | 0.053 | 20.000 | −8.465 | **<0.001** | −0.562 | −0.340 |

Significant p-values in bold, Bonferroni corrected for the number of ROIs ($p < 0.01$).
